# When is selection effective?

**DOI:** 10.1101/010934

**Authors:** Simon Gravel

**Affiliations:** McGill University, Montreal, Canada

## Abstract

Deleterious alleles can reach high frequency in small populations because of random fluctuations in allele frequency. This may lead, over time, to reduced average fitness. In that sense, selection is more ‘effective’ in larger populations. Recent studies have considered whether the different demographic histories across human populations have resulted in differences in the number, distribution, and severity of deleterious variants, leading to an animated debate. This article first seeks to clarify some terms of the debate by identifying differences in definitions and assumptions used in recent studies. We argue that variants of Morton, Crow and Muller’s ‘total mutational damage’ provide the soundest and most practical basis for such comparisons. Using simulations, analytical calculations, and 1000 Genomes data, we provide an intuitive and quantitative explanation for the observed similarity in genetic load across populations. We show that recent demography has likely modulated the effect of selection, and still affects it, but the net result of the accumulated differences is small. Direct observation of differential efficacy of selection for specific allele classes is nevertheless possible with contemporary datasets. By contrast, identifying average genome-wide differences in the efficacy of selection across populations will require many modelling assumptions, and is unlikely to provide much biological insight about human populations.

---

One of the best-known predictions of population genetics is that smaller populations harbor less diversity at any one time but accumulate a higher number of deleterious variants over time [1]. Considerable subsequent theoretical effort has been devoted to the study of fitness differences at equilibrium in populations of different sizes (e.g., [2]) and in subdivided populations (e.g., [3, 4]). The reduction in diversity has been observed in human populations that have undergone strong population bottlenecks: For example, heterozygosity decreased in populations that left Africa, and further decreased with successive founder events [5, 6, 7, 8]. The effect of demography on the accumulation of deleterious variation has been more elusive in both humans and non-human species. In conservation genetics, where fitness can be measured directly and effective population sizes are small, a modest correlation between population size and fitness was observed [9]. In humans, the first estimates of the fitness cost of deleterious mutations were obtained through the analysis of census data [10], but recent studies have focused on bioinformatic prediction using genomic data [11, 12]. Lohmueller et al. [13] found that sites variable among Europeans were more likely to be deleterious than sites variable among African-Americans, and attributed the finding to a reduced efficacy of selection in Europeans because of the Out-of-Africa (OOA) bottleneck. However, recent studies [14, 15] suggest that there has not been enough time for substantial differences in fitness to accumulate in these populations, at least under an additive model of dominance. By contrast Peischl et al. [16], and more recently Henn et al. [17], have claimed significant differences among populations under range expansion models, and Fu et al. [18] claims a slight excess in the number of deleterious alleles in European-Americans compared to that in African-Americans. These apparent contradictions have sparked a heated debate as to whether the efficacy of selection has indeed been different across human populations [19, 18]. Part of the apparent discrepancy stems for disagreement about how we should measure the effect of selection.

What does it mean for selection to be ‘effective’? Some genetic variants increase the expected number of offspring by carriers. As a result, these variants tend to increase in frequency in the population. This correlation between the fitness of a variant and its fate in the population—that is, *natural selection*—holds independently of the biology and the history of the population. However, the rate at which deleterious alleles are removed from a population depends on mutation, dominance, linkage, and demography, and can vary across populations. Multiple metrics have been proposed to quantify the action of selection in human populations and verify the classical population genetic predictions, leading to apparent discrepancies between studies.

In this article, we first review different metrics used in recent empirical work to quantify the action of selection in human populations. We show that many commonly used metrics implicitly rely on ‘steady-state’ or ‘equilibrium’ assumptions, wherein genetic diversity within populations is independent of time. This condition is not met in human populations. We discuss two measures for the efficacy of selection that are appropriate for the study of human populations and other out-of-equilibrium populations.

We then seek to provide an intuitive but quantitative understanding of the effect of mutation, selection, and drift on the efficacy of selection in out-of-equilibrium populations. This is done through a combination of extensive simulation and analytical work describing differentiation between populations after a split from a common ancestor. Using this information, we discuss how the classical predictions concerning the effect of demography on selection could be verified in empirical data from human populations.

## 1 Measuring selection in out-of-equilibrium populations

We consider large panmictic populations whose size *N(t)* may change over time, and whose reproduction follows the Wright-Fisher model [20]. Given alleles *a* and *A*, we assume that genotype *aa* has fitness 1, *aA* has fitness 1 + *s_i_ h_i_*, and *AA* has fitness 1 + s_i_. We suppose that *A* is the least favourable allele (*s_i_* < 0) and that 0 ≤ *h_i_* ≤ 1. In a random-mating population, an allele *A* at frequency *x_i_* adds an average of *δω_i_* = *s_i_* (2*h_i_ x_i_* + (1 – 2*h_i_*)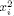) to individual fitness compared to the optimal genotype. We compute the expected fitness over multiple loci as 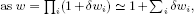 under the assumption that the individual selection coefficients *s_i_* are small. Finally, we define the genetic load *L* = 1 – *ω* as the total relative fitness reduction compared to the optimal genotype, *ω_max_* = 1. This yields

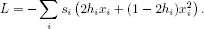

To study the effect of selection over short time spans and in out-of-equilibrium populations, we want to define instantaneous measures of the effect of selection on the genetic load and the frequency of deleterious alleles. In this article, the *rate of adaptation* refers the instantaneous rate of fitness increase (or load decrease) in a population. It has contributions from selection, mutation, and drift. The contribution of selection has been the object of considerable theoretical attention: It is the object of the Fitness Increase Theorem (FIT) (see, e.g., [20]). We will refer to the contribution of selection to the rate of adaptation as the *FIT efficacy of selection.*

We also wish to study the effect of selection on the *frequency* of deleterious alleles. There are multiple ways to combine frequencies across loci to obtain a single, genome-wide metric: Any linear function 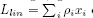 of the allele frequencies, with *ρ_i_* > 0 a weight assigned to locus *i*, provides an equally acceptable metric. A natural option, which weights alleles according to their selection coefficient, is Morton, Crow and Muller’s *total mutational damage* [21], which is equivalent to the *additive genetic load* that would be observed if all dominance coefficients were replaced by 1/2, i.e., 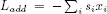. Mutation and selection systematically affect *L_lin_*, but genetic drift does not. We define the *Morton efficacy of selection* as the contribution of selection to 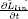. In simulations, where all alleles have equal fitness, we use *ρ_i_* = –*s*. Another common choice, in empirical studies, is to set *ρ_i_* = 1 for all sites annotated as deleterious by a prediction algorithm, and zero otherwise [15, 14, 18]. Since different empirical studies use different *ρ_i_*, direct comparison of the results can be challenging.

Because Morton and FIT efficacies are instantaneous measures of the effect of selection, they can be integrated over time to measure the effect of selection over arbitrary periods. Their integrals over long periods are directly related to classical steady-state metrics such as the rate of fixation of deleterious alleles and the average genetic load in a population.

To understand how genetic drift affects the FIT and Morton efficacy of selection, consider an allele with parental frequency *x*, selection coefficient |*s*| ≪ 1, and no dominance (*h* = 0.5). In the descending population, this allele is drawn with probability *x’* ⋍ *x* + *sx*(1 – *x*)/2. Figure 1 shows the resulting distribution in offspring allele frequency for *x* = 0.5, *s* = –0.6, and 2*N* = 100,500, and ∞. The average frequency *x’* is independent of *N*, hence the expected FIT and Morton efficacies are equal in all populations: Genetic drift does not instantaneously change the effect of selection.

**Figure 1.**
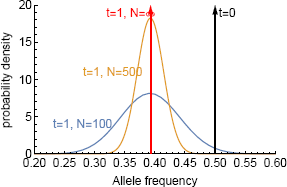
Frequency distribution of deleterious al-leles with initial frequency 0.5 after one generation, assuming population sizes of 2*N* = 100, 500, and ∞; a selection coefficient *s* = –0.6; and no dominance (*h* = 0.5). The average effect of selection after one generation does not depend on *N*.

If we let these populations evolve further, however, we will eventually find that deleterious allele frequencies decrease more slowly in smaller populations. This is because natural selection acts on fitness *differences*, and therefore requires genetic variation. By dispersing allele frequencies and reducing diversity, genetic drift also reduces the subsequent effect of selection (see Figure 2). Drift accumulated during one generation can change the efficacy of selection for many future generations. Conversely, the current average efficacy of selection depends on the drift accumulated in many previous generations. This delay between the action of drift and its impact on selection can be ignored in steady-state populations but not in out-of-equilibrium populations. For this reason, measures of the effect of selection that have been developed for populations of constant size can be misleading or biased when applied to populations out of equilibrium.

**Figure 2.**
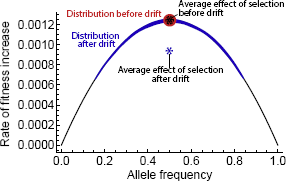
Effect of drift on the rate of adaptation for *h* = 0.5. We consider two populations with initial allele frequency 0.5 and selection coefficient *s* = –0.1. The red population does not undergo drift, and the blue population undergoes one generation of neutral drift, leading to increased variance in allele frequency. The reduced diversity at each locus leads to a lower average *rate* of fitness increase per generation (blue star). Because the rate of fitness increase is a concave function of allele frequency, drift always reduces the future effect of selection when *h* = 0.5.

### 1.1 Other measures of the effect of selection

The rate at which deleterious mutations are eradicated from a population, for example, is an intuitive metric for the effect of selection that has been recently applied to out-of-equilibrium populations [22]. Over long time scales or in the steady state, this rate of eradication is indeed equivalent to Morton’s efficacy of selection. However, in out-of-equilibrium populations, the rate of eradication is a biased measure of the effect of selection. In Figure 1, the smaller population has a higher rate of eradication of deleterious alleles, but this reflects the action of drift rather than the effect of selection. This effect of drift on the rate of eradication of deleterious alleles is short-lived on phylogenetic time scales, but it can be the dominant effect for time-scales relevant to human populations.

Classical work on the efficacy of selection in steady-state populations has emphasized the role of the combined parameter *Ns* in the dynamics of deleterious alleles. The importance of this combined parameter has led some authors to argue that it should be used as a metric for the efficacy of selection even outside the steady-state [15, 19]. This is problematic for practical and fundamental reasons. On the practical side, the parameter *N(t)s* is a function of time and does not allow for comparison between populations over finite times: *N(t)s* is not a rate, and its time integral is meaningless. At a more fundamental level, an instantaneous difference between two populations in the product *N(t)s* simply indicates a difference in effective population sizes. The interesting biological question is not whether the population sizes are different, but whether these differences lead to differential action of selection by the process illustrated in Figure 2.

More generally, it is commonly proposed that the effect of selection should be measured relative to the effect of drift [19], because the classical parameter *Ns* is a ratio between a selection term *s* and a drift term *1/N.* Such a relative measure is not necessary: Morton and FIT efficacies are absolute measures of the effect of selection and they do capture the classical interaction between selection and genetic drift: In populations of constant size, these efficacies do depend on the relative magnitude of selection and drift *coefficients* through the classical parameter *Ns.* In out-of equilibrium populations, however, they depend on a more complex function of *s* and *N(t).* In other words, the classical parameter *Ns* does not measure the effect of selection *as compared to* the effect of drift; but rather the effect of selection *as modulated by* past genetic drift.

Finally, even though most classical work has focused on the effect of selection on fitness or allele frequency, Henn *et al.* [17] recently proposed to measure the effect of selection on *diversity*, defining a ‘reduction in heterozygosity’ *(RH)* statistic that compares the heterozygosity of selected and neutral sites. We show in Section S1.2 that *RH* is robust to the effect of genetic drift, but it can be biased by recent mutations.

## 2 Asymptotic results

To study the effect of selection after a population split, we calculate the moments of the expected allele frequency distribution *φ(x, t)* under the diffusion approximation. In this formulation, *ϕ(x,t)dx* represents the expected number of alleles with frequency between *x* and *x* + *dx* at time *t*. In a randomly mating population of size *N* = *N(t)* ≫ 1 and constant *s* and *h*, the evolution of *ϕ(x, t)* approximately follows the diffusion equation [20]:

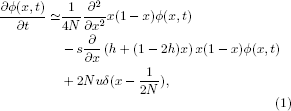

where *u* is the total mutation rate. The first term describes the effect of drift; the second term, the effect of selection; and the third term describes the influx of new mutations: *δ* is Dirac’s delta distribution. From this equation, we can easily calculate evolution equations for moments of the expected allele frequency distribution *µ_k_* = 〉*x^k^*〉. For example, the rate of change in allele frequencies 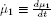, is driven by mutation and selection:

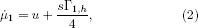

where Γ*_i, h_* = 4(*µ_i_* – *µ*_*i*+1_)*h* + 4(1 – 2*h*)(*µ*_*i*+_1 – *µ*_*i*+2_) is a function of the diversity in the population that generalizes the heterozygosity π_1_ = Γ_1,1/2_ (see Appendix for detailed calculations, and References [23, 24] for other applications of the moment approach). We can define the contributions of selection and mutation to changes in allele frequency as 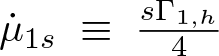 and 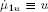. Morton’s efficacy of selection at a locus is simply 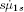. Whereas the effect of mutation is constant and independent of population size, Morton’s efficacy depends on the history of the population through Γ_1, *h*_:

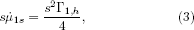

Similarly, changes in the expected fitness *W* can be decomposed into contributions from mutation, drift, and selection:

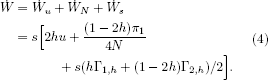

Favorable mutations increase fitness, drift increases fitness when fitness of the heterozygote is below the mean of the homozygotes, and selection always increases average fitness.

The FIT efficacy, 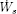, is therefore

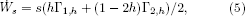

The right-hand side is the additive variance in fitness, and Equation (5) is an expression of the Fitness Increase Theorem (see, e.g., Equations 1.9 and 1.42 in [20]). Importantly, the FIT efficacy only describes one of three genetic contributions to the rate of adaptation. Interpreting changes in fitness in terms of FIT efficacy requires picking apart the effects of drift and mutation from those of selection. In addition to these genetic effects, changes in the environment can directly affect fitness, introducing a further confounder [25].

Now consider an ancestral population that splits into two isolated randomly mating populations with initial sizes *N_A_* and *N_B_* at time *t* = 0. The populations may experience continuous population size fluctuations. If we expand the moments *µ_k_* of the allele frequency distribution in Taylor series around *t* = 0, we can easily solve the diffusion equation to study the differentiation between the two populations right after the bottleneck. Here we provide an overview of the main results. Detailed derivations are provided in the Appendix.

The difference in fitness between the two populations, Δ*W*(*t*) = *W_A_* (*t*) – *W_B_* (*t*) grows linearly in time under dominance:

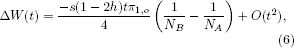

Here, *t* is measured in generations, π_1,*o*_ is the expected heterozygosity in the source population, and *O*(*t*^2^) represents terms at least quadratic in *t*. This rapid, linear differentiation is driven by drift coupled with dominance. The smaller population has higher fitness when *h* > 0.5 for *s* < 0: Drift hides dominant, deleterious alleles from the action of selection.

If the source population is large and *h* > 0, we have 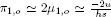 [26] and the rate of fitness differentiation is independent of *s*. This generalizes Haldane’s observation that load is insensitive to the selection coefficient in large populations [27]. By contrast to the constant-size population case, however, the observation does not hold when *h* = 0. The initial response to the bottleneck is independent of fitness for 0 < *h* < .5 (see Figures S2 and S3), but not for *h* = 0 or *h* = 0.5 (see Figures 3 and S1).

**Figure 3.**
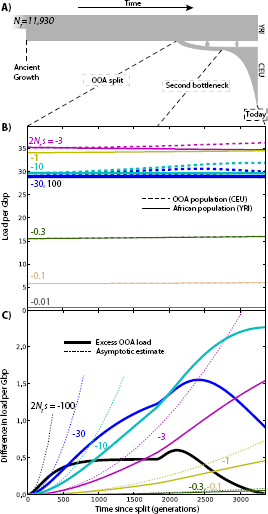
Changes in load after the OOA split model illustrated in (A) as a function of s, given ancestral population size *N_r_* = 11, 930 and no dominance (*h* = 0.5). (B) Load evolution after the split. We subtracted the load due to variants fixed in all populations. (C) Difference in load between populations. Since *h* = 0.5, this is equivalent to the difference in additive load. Dotted lines show asymptotic results from Equations (8) and (10). Load is given per Gbp of variants at the specified selection coefficient. The total amount of variation under strong selection in the human genome is likely much less than 1 Gbp.

The effect of selection on fitness differences, Δ*W_s_*(*t*), grows only quadratically:

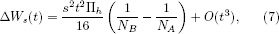

where Π_*h*_ is a measure of diversity that reduces to π_1, *o*_ when *h* = 0.5 (see Appendix). This slower response is the mathematical consequence of the intuition provided by Figures 1 and 2: Right after the split, the fitnesses are identical and the efficacy of selection is the same in both populations. It takes time for drift to increase the variance in allele frequency and cause differences in the efficacy of selection, accounting for a factor 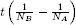. It then takes time for differences in the efficacy of selection to accumulate and produce differences in fitness, accounting for an additional factor *st*.

Combining Equations (6) and (7), we get an asymptotic result for the load differentiation.

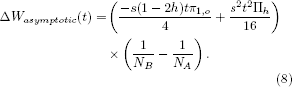

This expression describes the leading differentiation in fitness in all simulations below. It is straightforward to refine this asymptotic result by computing higher-order corrections, however the number of terms in the expansion increases rapidly. Some of these terms are of particular interest, such as the contribution of new mutations. Since the direct effect of mutation on load is independent of demography [Equation (4)], we must wait for mutations to accumulate before load differentiation can begin. This leads an additional factor of *ut* compared to the case of standing variation. The contribution of drift acting on new recessive mutations is therefore quadratic:

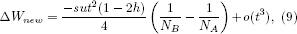

The effect of selection on new mutations is only cubic in time: We must wait for mutations to appear (contributing a factor of *ut*), then wait for drift to cause differences on the frequency distribution of the new mutations [contributing a factor of 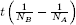], and finally wait for selection to act on these frequency distribution differences (contributing a factor of *st.)* The leading contribution of selection is therefore

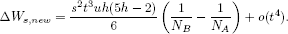

Finally, since drift alone does not produce differences in average allele frequencies, the rate of differentiation in deleterious allele frequencies is always quadratic in time:

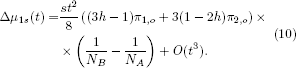

## 3 Simulations

We simulated the evolution of *ϕ(x, t)* using ∂a∂i [28] and the Out-Of-Africa (OOA) demographic model illustrated in Figure 3A. This model begins with an ancestral population of size *N_r_* = 11930 with frequency distribution following the quasi-stationary distribution of Kimura [29], and features population splits and size changes that were inferred from synonymous polymorphism from the 1000 Genomes dataset [30]. We estimated the probability *ϕ_n_(i, t)* that a variant is at frequency i in a finite sample of size *n* = 100 for each population, given a mutation rate of *µ* = 1.44 × 10^−8^bp^−1^ generation^−1^ [31] in an infinite genome. We used the finite sample predictions to estimate the expected genetic load and the contributions of drift, selection, and mutation. Finally, to ensure that results were not model-dependent, we repeated each simulation using a different demographic model described in [15], featuring a single deeper but shorter OOA bottleneck.

We simulated all combinations of selection coefficients 2*N_r_s* ∈ {0, –0.01, –0.1, –0.3, –1, –3, –10, –30, –100}, and dominance coefficients *h* ∈ {0,0.05,0.2,0.3,0.5,1}. The contributions of selection and drift were obtained using Equation (4). To emphasize the long-term effects of the OOA bottleneck even after drift is suppressed, simulations were also carried to future times assuming large population sizes (*N* = 20*N_r_*) and no migrations (Figure S6). In all cases, Equations (8), (9), and (10) capture the initial increase in load (Figures 3, S1, S2, and S7).

### 3.1 Genic selection; *h* = 1/2

Simulated differences in load are modest and limited to intermediate-effect variants (.3 < |2*N_r_s*| < 30, 2.5x10^−5^ < |*s*| < 0.0025). Assuming the distribution of fitness effects inferred from European-American data by Boyko *et al*. [32], the excess load in the OOA population is 0.49 per Gb of amino-acid-changing variants, in addition to a total accumulated load of 24 per Gb in the African population (this accumulated load does not include variation that was fixed at the time of the split). If we consider the 24 Mb of exome covered by the 1000 Genomes project, and assume that 70% of mutations are coding in that region [33], the model predicts a non-synonymous load difference of 0.008. The total estimated non-synonymous load, excluding mutations fixed in the ancestral state, is 0.4 in the African-American population. In this model, the reduced efficacy of selection caused by the OOA bottleneck leads to a relative increase in non-recessive load of 2%. Since we did not consider fixed ancestral deleterious alleles in the total load, this figure is an overestimate of the relative increase in load due to the bottleneck. The relative increase reaches a maximum of 8% for mutations with –20 < 2*N_r_s* < –10. The results are similar if we use the distribution of fitness effects inferred from African-American data [32].

Using the simple bottleneck demographic model of Do et al [15], we find very similar load (24 per Gbp) and load differences across populations (2% of the total load).

### 3.2 Partial and complete dominance

The picture changes dramatically when we consider recessive deleterious variants (*h* = 0). Reactions to changes in population size are linear rather than quadratic, and they are more substantial than in the additive case (Figure S1). The OOA load due to segregating variants with 2*N_r_s* = –100 almost doubles after 500 generations. This excess load in the OOA population is due entirely to drift, and leads to an increased efficacy of selection in the OOA population, since a higher proportion of deleterious alleles are now visible to selection. The difference in load for the most deleterious variants is therefore not sustained.

Both the number of very deleterious variants and the associated genetic load eventually becomes higher in the simulated Yoruba population. By contrast, weak-effect deleterious variants contribute more load in the simulated European population.

Even though a bottleneck inexorably leads to increased load when no dominance is present, the additional exposure of recessive variants therefore leads to ‘purging’, a reduction of the frequency of deleterious alleles (see [2] and references therein). Simulations show that the increase in recessive load can last hundreds or thousands of generations for weakly deleterious variants. Gl´emin argued that the purging effect is suppressed in constant-sized population when *Ns* is much less than “2 to 5”. [2]. This also holds in a non-equilibrium setting in recessive alleles going through a bottleneck (Figure S1, see also [34]). The time required for purging to compensate the initial fitness loss increases rapidly as the magnitude of the selection coefficients decreases: Whereas our model predicts a reduced load in present-day OOA populations for alleles with 2*N_r_s* = –100, it would take over 20, 000 generations of continued isolation in large constant-sized population to see purging in alleles with 2*N_r_s* = –3 (Figure S6).

Opposite effects are observed for dominant deleterious variants (Figure S7). Drift tends to increase fitness by combining more of the deleterious alleles into homozygotes, reducing their average effect on fitness. The difference between populations is much less pronounced and less sustained than in the recessive case. Equation (6) shows that the reduced magnitude is caused by reduced ancestral heterozygosity, π_1, 0_: Dominant deleterious alleles are much less likely to reach appreciable allele frequencies before the split. Here again, the population with the highest load depends on the selection coefficient, with a higher load in the simulated European population for strongly deleterious variants and a higher load in the simulated Yoruba population for the weakly deleterious variants.

The distribution of dominance coefficients for mutations in humans is largely unknown, but non-human studies suggest that partial recessive may be the norm (see, e.g., [35] and references therein). Under models with *h* = 0.2, we find that the genetic load is elevated in OOA populations for most selection coefficients *N_r_s*, whereas the *additive* genetic load is mostly reduced (Figure S3B-C and S4B-C). These simulations suggest that the rate of adaptation was reduced in OOA populations (i.e., the genetic load is higher in OOA population), while the efficacy of selection was higher in the OOA population, whether it is measured by the Morton efficacy or the FIT efficacy (Figure S5). Thus, unless most nearly-neutral variation has *h* > 0.20, we do not expect an overall elevated number of deleterious variants in OOA populations. As we move closer to additive selection, for example at *h* = 0.3, the contributions of alleles with larger and weaker selection coefficient are of comparable magnitude and opposite direction. Because of our limited ability to estimate selection coefficients in humans, this might explain why observing differences in the overall frequency of deleterious alleles between populations has been so difficult. This also suggests that any claim for an across-the-board difference in the efficacy of selection between two populations will have to rely on a number of assumptions about fitness coefficients in human populations.

## 4 Present-day differences in the efficacy and intensity of selection

The Wright-Fisher predictions for the instantaneous Morton and FIT efficacies of selection, Equations (3) and (5), depend on the present-day allele frequency distribution, on the dominance coefficient h, and on the selection coefficient s. However, s is a multiplicative factor in both equations and cancels out when we consider the relative rate of adaptation across populations. We can therefore use Equations (3) and (5) to estimate differences in the efficacy of selection between populations based on the present-day distribution of allele frequencies. For nearly-neutral alleles, the present-day frequency distribution is similar to the neutral frequency spectrum and largely independent of h. We can therefore use the present-day frequency spectrum for synonymous variation to estimate the relative efficacy of selection for all nearly-neutral alleles at different values of *h* (Figure 4). Figures S10 and S11 show similar results for non-synonymous and predicted deleterious alleles (For the most deleterious classes, the assumption that the present-day frequency spectra depend weakly on *h* is less accurate).

In the nearly-neutral case, the Luhya population (LWK) shows the highest Morton an FIT efficacy of selection for most dominance parameters and is used as a basis of comparison. The estimated FIT efficacy of selection is higher in African population for all dominance coefficients, as is the Morton efficacy, except for completely recessive alleles. The reduction in Morton’s efficacy of selection for nearly-neutral variation in OOA populations is 25% to 39% for dominant variants, 19% to 31% for additive variants, and 6% to 13% for fully recessive variants. The reduction in the FIT efficacy in OOA populations is 29% to 44% for dominant variants, 19% to 31% for additive variants, and 0.2% to 6% for fully recessive variants. This is also consistent with the interpretation of Gl´emin that purging, the reduction in the frequency of recessive alleles caused by a bottleneck, is not expected for nearly neutral variants. By contrast, estimates using sites with high predicted pathogenicity according to CADD [36] do suggest that purging of deleterious variation by drift is still ongoing in OOA populations (Figures S10 and S11).

Admixed populations from the Americas with the highest African ancestry proportion also show elevated efficacy of selection: African-Americans (75.9% African ancestry [37]), Puerto Rican (14.8% African ancestry [31]), Colombians (7.8% African ancestry [31]), and Mexican-Americans (5.4% African ancestry [31]). The Morton efficacy of selection in admixed populations is much larger than the weighted average of source populations would suggest (Figure 4C, which uses CHB, CEU, and YRI as ancestral population proxies for Native, European, and African ancestries). By averaging out some of the genetic drift experienced by the source populations since their divergence, admixture increases the overall amount of additive variance in the population, and therefore leads to a substantial and rapid increase in the predicted efficacy of selection for nearly neutral alleles.

**Figure 4.**
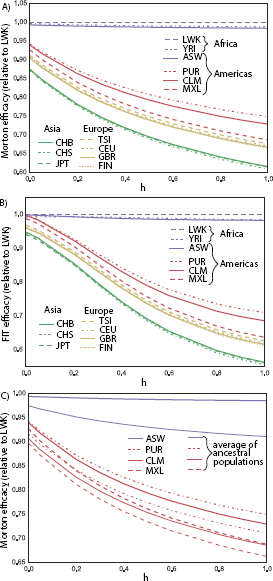
A) Present-day Morton efficacy of selection for nearly neutral variants, estimated from 1000 Genomes synonymous variation. B) FIT efficacy for the same variants, and C) Morton efficacy in admixed populations is increased compared to the average efficacies in ancestral populations.

## 5 Discussion

Selection affects evolution in many ways. It tends to increase the frequency of favourable alleles and the overall fitness of a population, and it often reduces diversity. The rates at which it performs these tasks varies across populations, and population geneticists like to frame these differences in terms of the efficacy of selection. The word ‘efficacy’ implies a measure of achievement, but there are many ways to define achievement for selection. We considered two measure of achievement: the change in deleterious allele frequency (i.e., Morton’s efficacy), and the change in load caused by selection (i.e., the FIT efficacy). Even though the two quantities are closely related, and are equal for additive selection, Morton’s efficacy is much easier to measure: systematic differences in the frequency of deleterious alleles are robust to drift and to modest changes in the environment. By contrast, the FIT efficacy is impossible to observe directly and requires picking apart the contributions of selection, drift, and the environment. Given the long-standing controversy about how this should be done in the context of Fisher’s Fundamental Theorem [38], we would advise against using it.

We have argued that other popular measures for the efficacy of selection [13, 19, 8, 17] are biased in out-of-equilibrium populations studied over short time-scales. Many previous claims that selection acted differentially in human populations [13, 8] could be explained by these biases. Confirming this interpretation, Fu et al. [18] found no differences in the average frequency of deleterious alleles between African-Americans and European-Americans in the ESP 6500 dataset [39]. However, they did report a slight but extremely significant difference in the average number of deleterious alleles per individual for a set of putatively deleterious SNPs. The contrasting results are surprising, since the two statistics are equal up to a multiplicative constant: the average number of deleterious alleles per genome equals the mean frequency of deleterious alleles multiplied by the number of loci. We could reproduce the results from [18], but found that the statistical test used did not account for variability introduced by genetic drift in a finite genome: results remained significant if allele frequencies were randomly permuted between African-Americans and European-Americans (see Section S1.1 for details). This emphasizes that an empirical observation of differences in genetic load must be robust to both finite sample size and finite genome to be attributed to differences in the efficacy of selection.

Figures 4, S10 and S11 strongly suggest that the OOA bottleneck still influences the present-day efficacy of selection. By extension, they also suggest that the efficacy of selection did differ and will differ among populations. Importantly, the differences in frequency distributions across populations that provide this support are not a *consequence* of past differences in the efficacy of selection but a possible *cause* for such differences in the present and future. We have shown that some of the future differences are not inevitable and can be attenuated by demographic processes including admixture. Therefore, measuring actual differences in the efficacy of selection can only be achieved by measuring actual differences in the average frequency or effect of deleterious alleles.

Simulations presented here, together with the results of [14, 15], do suggest that the classical prediction on the differential efficacy of selection in small populations can be verified if only we can accurately isolate variants of specific selective effect and dominance coefficients. By picking apart variants of different selection and dominance coefficients, we should soon be able to convincingly and directly observe the consequences of differences in the efficacy of selection. The recent results of Henn et al. [17], using a version of Morton’s efficacy, do suggest such differences for a subset of variants and therefore provide important experimental validation for a classical population genetics prediction. By contrast, the observation of genome-wide differences in the efficacy of selection across populations depends on the cancellation of effects across different variant classes, and can therefore depend sensitively on the particular choice of a metric. For this reason, overall differences in load among populations may not be particularly informative about the fundamental processes governing human evolution.

## 6 Acknowledgements

I thank S. Baharian, M. Barakatt, B. Henn, D. Nelson, and S. Lessard for useful comments on this manuscript, and W. Fu and J. Akey for help in reproducing their results. This research was undertaken, in part, thanks to funding from the Canada Research Chairs program and a Sloan Research Fellowship.

## A1 Appendix

### A1.1 Background

To derive the asymptotic results in the text, we start with the diffusion approximation for the distribution *ϕ(x, t)* of allele frequencies *x* over time *t* in an infinite-sites model (see [26], section 8.6):

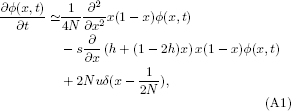

where *N* is the effective population size, *h* is the dominance coefficient, *s* is the selection coefficient, and *u* is the mutation rate. In this model, new mutations are constantly added via Dirac’s delta function *δ*. Because there are no back mutations in this model, the proportion of fixed mutations increases over time without bound. Because we are only interested in population differences accumulating over a short time span, however, we can simply ignore the (infinite) number of deleterious alleles that fixed before the population split. The time-scales that we will consider are short enough that back-mutations and multiple mutations contribute little to changes in allele frequencies.

A complete solution of this problem can be expressed as a superposition of Gegenbauer polynomials [29]. However, here we are looking for simple asymptotic results that will help us understand the dynamics of the problem. We will consider the evolution of moments of the allele frequency distribution: 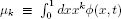. Similar moment approaches have been used in [23, 24]. Because there is a possibly infinite number of fixed sites at frequencies 0 and 1, it is often convenient to distinguish contributions from segregating sites and fixed sites:

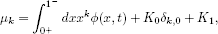

where *K_0_* is the number of sites fixed at frequency 0, *K_1_* is the number of sites fixed at frequency 1, and *δ*_*k*, 0_ is Kronecker’s delta. Both *K_0_* and *K_1_* can be infinite in this model, but this will not be a problem since we will ultimately consider only differences or rates of change in the moments, and these remain finite. In this notation, *µ*_0_ is the (possibly infinite) number of sites, and *µ*_1_ is the expected number of alternate alleles per haploid genome.

To obtain evolution equations for the moments, we integrate both sides of equation (A1) using 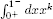. The left-hand side gives

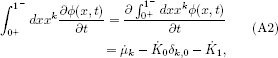

and the right-hand side can be integrated by parts. For *k* = 0, this yields

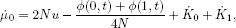

where *ϕ*(0, *t*) and *ϕ*(1, *t*) are defined by continuity from the open interval (0,1) and do not include fixed sites. Because the number of sites is constant 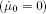 and the diffusion equation is continuous, we require

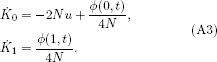

These equations are equivalent to Equations 3.18 and 3.19 in [29].

To obtain an evolution equation for *µ_k_* at arbitrary *k*, we return to the integration of Equation (A1) with 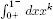. We use the left-hand-side expression obtained in Equation (A2), and once again integrate the right-hand side by parts. This yields

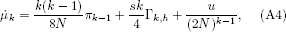

where

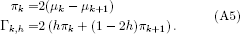

These are functions of the moments *µ* and can therefore be thought of as measures of the shape of the frequency distribution *ϕ*.

The first term in (A4) represents the effect of drift, the second term the effect of selection, and the third term the effect of mutation.

For example, if *k* = 1 and *h* = 1/2, we get

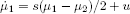

The frequency of damaging alleles can decrease because of selection, or increase because of mutation.

### A1.2 Response in allele frequencies

Solving Equation (A4) in general is challenging, because 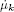 can depend on *µ*_*k*+1_ and *µ*_*k*+2_, leading to an infinite number of coupled equations. However, it can be used to calculate the response in allele frequency to a sudden change in demographic or selective conditions. Consider a population of size *N_o_* that experiences a change in size to *N_A_* at time *t* = 0. We can expand *µ_k_* for short times:

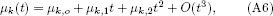

where *µ*_*k*, o_ is the kth moment prior to the population size change and *O*(*t*^3^) represent terms at least cubic in *t*. The coefficients can be evaluated by expanding both sides of Equation (A4) using Equation (A6), then collecting powers of *t*. For example, we get

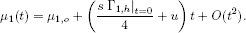

The frequency of variants can increase even in a steady-state regime with *N_A_* = *N_o_*, since our model assumes a constant supply of irreversible mutations. However, this linear term is independent of *N_A_* and does not contribute to differences across populations that share a common ancestor. Differences Δ*µ*_1_(*t*) in the number of segregating sites between two populations with sizes *N_A_* and *N_B_* appear at the next order in *t*. Computations are elementary but a bit more cumbersome. Matching terms linear in *t* in Equation (A4), we find equation (10):

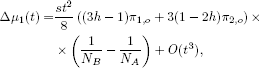

where π_*i, o*_ is the moment π_*i*_ computed for the common ancestral population.

### A1.3 Response in genetic load

To compute the fitness in the diploid case, we write

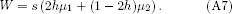

Using (A4), we get

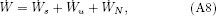

where

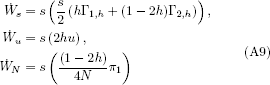

are the instantaneous contributions of selection, mutation, and drift to changes in fitness. The mutation term is constant in time and independent of population size; it does not directly contribute to differences across populations. The drift term, by contrast, has an explicit dependence on the population size; this leads to differentiation between populations that grows linearly in time. To see this, we compute the load using Equation (A7) and the time dependence computed in Equation (A6), as in section A1.2:

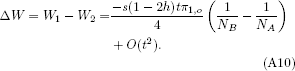

This reduction in load is driven by drift, i.e., the third term in equation (A9). It is not caused by selection, in the sense that it does not result from differential reproductive success between individuals. As expected, the contribution of drift vanishes for additive variants (*h* = 1/2).

For arbitrary *h*, the change in fitness caused by selection is

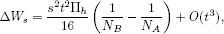

where Π_*h*_ is a statistic of the ancestral frequency distribution:

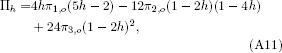

which reduces to the heterozygosity π_1,0_ when *h* = 1/2. The statistic Π_*h*_ depends only on the ancestral frequency distribution and the dominance coefficient.

Genetic drift also contributes to the changes in load at second order in *t* through 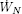. In addition to the linear term from Equation (A10), we find three quadratic contributions that vanish when *h* = 1/2: a second-order contribution of genetic drift, a contribution from the rate of change in population size and drift, and a contribution from new mutations and drift. Even though these terms can be comparable in magnitude to the contribution of selection in Equation (A11) when *h* ≠ 1/2, they are sub-dominant to Equation (A10). We only consider the contribution of new mutations in some detail, as this contribution tells us whether population differentiation in the genetic load is due to old, shared variation or to new, population specific variation.

### A1.4 Effect of new mutations

If we set π_*i,o*_ = 0 in the equations above, we can calculate the impact of new mutations on the genetic load. The leading term is again due to drift and dominance:

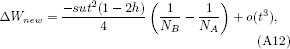

while the leading term describing the efficacy of selection is now cubic in *t:*

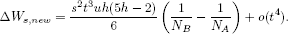

When *h* ≠ 1/2, drift also contributes *t^3^* terms to *ΔW_new_*. These are reasonably straightforward to compute, but are sub-dominant to Equation (A12).

We therefore use the asymptotic result:

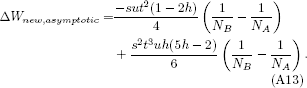

Comparisons with simulated data are shown on Figure S9.

### 1 Supplementary Material

#### S1.1 Analysis of the Fu et al. result

Fu et al presents two different tests for differences in the number of deleterious alleles between individuals. In the first test, a t-test, they compare the average frequency of deleterious alleles between the two populations and find no significant differences (*p* = 0.82). In the second test (a Mann-Whitney test), they compared the number of deleterious alleles per individual, and found an extremely significant difference (*p* < 10^−15^). We could reproduce these results using the publicly available ESP 6500 data [39] using a permutation test. Since the average frequency of deleterious alleles and the number of deleterious alleles per individual are related by a proportionality constant (i.e., the length of the genome), we suspected that the significant difference results from a difference in the underlying null models of each test.

To verify this, we produced simulated datasets in which the allele frequency at each site was randomly permuted between European-Americans and African-Americans, creating a dataset for which there is no meaningful evolutionary difference between the two populations, except for randomly assigned differences in allele frequency. We nevertheless found very significant differences between the average number of deleterious alleles carried by individuals of the two populations for most simulations. This effect can even be reproduced by analyzing a single SNP. This is the Mann-Whitney test used in Fu et al. conditions on the set of SNPs used: it does not account for the fact that drift can have affected different SNPs differently. The difference in deleterious allele count observed in Fu et al is therefore real, but it only applies to a particular set of SNPs and it is consistent with the action of genetic drift acting on neutral variation. It does not indicate differences in the action of selection, nor systematic differences in fitness across populations. To assign differences in deleterious allele frequency to the systematic action of selection, one must show that the difference is robust to both finite sample size and finite genome effects. Both can be tested through simple resampling strategies. In the ESP6500 example, resampling over SNPs by bootstrap led to no significant differences in the number of deleterious alleles per individual.

#### S1.2 Reduction in heterozygosity (RH) statistic

The reduction in heterozygosity *(RH)* statistic was recently introduced [17] as a tool to measure the effect of selection on population diversity as an addition to analyses based on Morton’s efficacy. In this section, we show that even though *RH* is an interesting measure of diversity in a population, its interpretation in terms of the effect of selection can be challenging: First, *RH* can be affected by recent mutation and, second, differences in the effect of selection on *RH* may reflect rather mundane normalization issues rather than interesting biology.

*RH* is defined as

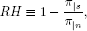

where π is the average heterozygosity, |*s* indicates a quantity measured at selected sites, and |*n* indicates a quantity measured at neutral sites. The rate of change in RH is therefore

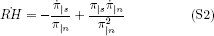

Using equation (A4) and (A5) to leading order in 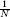, we find

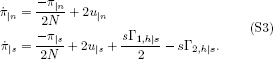

where *u* is the mutation rate and Γ_*i,h*_ is defined in Equation (A5). The first terms describe the action of drift, the second terms describe the action of mutation, and the last two terms in the second line describe the action of selection. The contributions of drift cancel out in Equation (S2), so that *RH* only depends on contributions from selection and mutation:

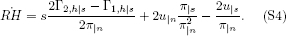

Now consider the rate of change in the difference *ΔRH* between populations *A* and *B:*

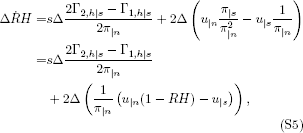

By contrast with Morton’s efficacy, the mutation term does not cancel out for *ΔRH*.

To show that the selection term can be substantial, we consider the case of strong selection, where 1 – *RH* ≪ 1. In this case, the contribution of mutation to 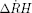 is

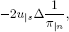

while the contribution of selection to 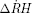 is

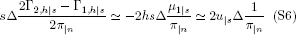

where the first approximation uses Equation (A5) and assumes that higher moments are sub-dominant under strong selection because allele frequencies are small. The second approximation uses the mutation-selection balance relation 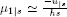 (e.g, equation 6.2.9 in [26]). The effects of mutation and selection on Δ*RH* cancel out: average frequencies of very deleterious alleles are governed by the mutation-selection balance and are unaffected by population size differences. Mutation tends to increase RH in the less diverse population.

Identifying the effect of selection on diversity therefore requires picking apart the effects of selection and mutation (if measuring Δ*RH*), or the effect of selection and drift (if measuring Δπ_*s*_). As in the case of fitness, knowing only the present-day distribution of allele frequency is not enough to identify the effect of selection unambiguously, unless the alternate effect can be shown to be weak.

#### S1.3 Example of two populations with identical start and end frequency distributions, but different measures of FIT efficacy

Imagine that all alleles all have dominance coefficient 0 < *h* < 1, selection coefficient *s* and dominance *h*, and consider two identical populations with exactly *L* segregating alleles, all at frequency *x_o_* at time *t_o_*. The initial fitness in both populations is *W_o_* = *Ls* (2*hx_o_* + (1 – 2*h*)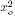). Assume no mutation. In the second population, a short bottleneck occurs that increases the variance in allele frequency before selection had time to change mean allele frequencies. The difference in fitness between the two populations after the short bottleneck is *δW*_bottleneck_ = *Ls*(1 – 2*h*)*σ*^2^, where *σ*^2^ is the variance in allele frequency in the second population. The change in fitness in population 2 up to this point is entirely attributed to drift.

After the bottleneck, the population sizes increase to a very large value, so that subsequent effects of drift can be neglected. Selection is left to act until we have approached the maximum fitness state, which has fitness *W* = *Ls* if *s* > 0 and *W* = 0 if *s* < 0. The change in fitness *W* – *W_o_* is equal in both populations, since the initial and final states are exactly the same. The FIT ‘effect of selection’ is *W* – *W_o_* in population 1, since all changes are due to selection. In population 2, the effect of selection is *W* – *W_o_* – *δW*_bottleneck_, because a change *δW*_bottleneck_ was caused by driftrather than by selection.

#### S1.4 Microscopic and macroscopic efficacy and intensity of selection

The main text discusses the effect of selection by averaging over all possible allelic trajectories. A natural question is whether individual allelic trajectories can be interpreted in a similar manner. Of course, it is not possible in general to attribute specific changes in frequency to the effect of drift or to selection, just as it is impossible to attribute the precise number of offspring borne by an individual to drift or fitness alone—these attributions can be made only in an average sense. However, individual trajectories can contain more information than allele frequency distributions, for example when selection coefficients fluctuate. Recent work has defined a microscopic ‘rate of adaptation’ [25] as a measure of the effect of selection on individual trajectories. Recent work has defined a microscopic ‘rate of adaptation’ [25] as a measure of the effect of selection on individual trajectories. Here we show how the different metrics of the impact of selection can be applied to individual trajectories, and how they relate to Mustonens rate of adaptation.

Consider an individual allelic trajectories. If the allele frequency has trajectory {*x*_t_}_*t*= 1,…, *T*_, with *t* the time in generations, we can write the fitness change Δ*W_t_* at generation *t* as

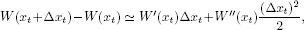

where Δ*x_t_* = *x*_*t*+1_ –*x_t_* and *W*′ represents the partial derivative of the fitness function with respect to frequency *x*. In the constant-fitness models discussed above, *W*′(*x_i_*) = 2*s*(*h* + (1 – 2*h*)*x*) and *W*″(*x_i_*) = 2*s*(1 – 2*h*).

The expectation of *W*′(*x_t_*)Δ*x_t_* gives our 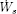, and the expectation of 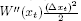 gives 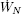 when |*s*| ≪ 1. We therefore define the quantities *σ_t_* = *W*′(*x_t_*)Δ*x_t_* and *ν_t_* = *W*″(*x_t_*)(Δ*x_t_*)^2^ as microscopic analogs of the macroscopic efficiency of selection 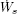 and drift 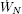. These are not the only possible analogs—for example, we could consider the expectation of the linear term 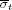 as the microscopic effect of selection, and 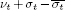 as the microscopic effect of drift, without changing the expected values.

Mustonen and Lassig define a ‘fitness flux’ *ϕ_t_* in [25] as a measure of the rate of adaptation. The fitness flux definition appears identical to our definition for *σ*, namely 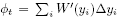, where *y_i_* is an allele frequency trajectory sampled densely in time. However, it is not equal but corresponds to the total change in fitness, *ν_t_* + *σ_t_*. Whereas our trajectory {*x*}_t_ is labeled by the time in generation, the time steps in {*y_i_*} are chosen so that 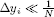. Because of this constraint, time steps in [25] must be finer than one generation, and the *y_i_* must be interpolated within generations. While integrating over this smoothed trajectory, quadratic terms in Δ*y_t_* can be neglected: The integrated fitness flux yields the total rate of fitness change, whether it is due to drift or selection.

The intensity of selection *i* measures the rate of increase in frequency of favorable alleles. At a microscopic level, the intensity of selection is simply

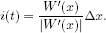

When 0 ≤ *h* ≤ 1 and selection coefficients do not change sign over time, 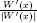 is 1 or favorable alleles, and -1 for deleterious ones: The intensity of selection is simply the average change in frequency of favorable alleles. If *h* < 0 or *h* > 1, we have overdominance, and the favored allele is frequency dependent. In that case, the intensity of selection is also independent of the trajectory followed, and is given by *I* = – sgn(*sh*) 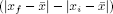, where 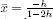 represents the frequency of optimal fitness. As discussed in the main text, there are many ways to weight the intensity of selection when the effect is to be measured across multiple sites. As long as the weighing scheme is not frequency or time-dependent, it remains possible to directly compare the intensity of selection across populations without the need for detailed modeling.

**Figure S1.**
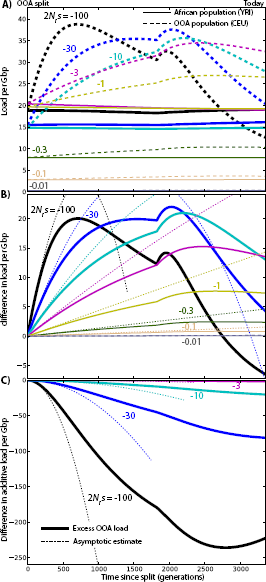
Changes in load after the OOA split model illustrated in Figure 3A, assuming a recessive variant (*h* = 0.0), as a function of s and given ancestral population size *N_r_* = 11, 930. (A) Overall genetic load evolution. The load due to variants that are fixed in all populations is not included. (B) Difference in load between the two populations. (C) Differences in additive load. The dashed lines represent the asymptotic results from Equations (8) and (10).

**Figure S2.**
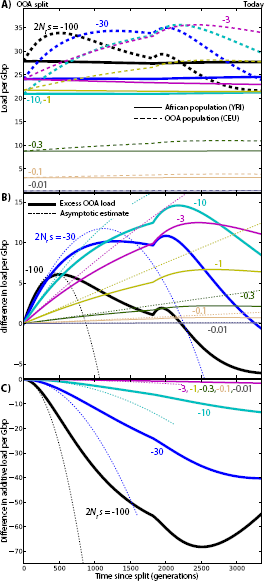
Changes in load after the OOA split model illustrated in Figure 3A, assuming partial recessive variants (*h* = 0.05) as a function of s and given ancestral population size *N_r_* = 11, 930. (A) Overall genetic load evolution. The load due to variants that are fixed in all populations is not included. (B) Difference in load between the two populations. (C) Differences in additive load. The dashed lines represent the asymptotic results from Equations (8) and (10).

**Figure S3.**
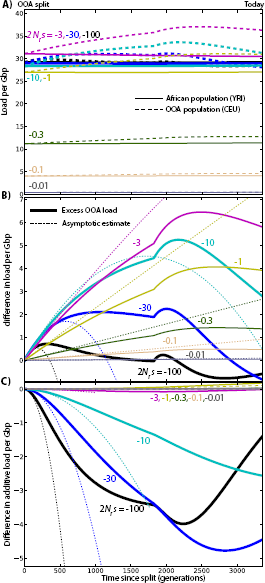
Changes in load after the OOA split model illustrated in Figure 3A, assuming partial recessive variants (*h* = 0.2) as a function of s and given ancestral population size *N_r_* = 11, 930. (A) Overall genetic load evolution. The load due to variants that are fixed in all populations is not included. (B) Difference in load between the two populations. (C) Differences in additive load. The dashed lines represent the asymptotic results from Equations (8) and (10).

**Figure S4.**
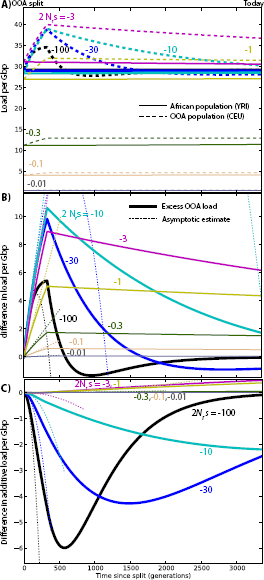
Changes in load after the simple OOA bottleneck model from [15], assuming partial recessive variants (*h* = 0.2) as a function of s and given ancestral population size *N_r_* = 11, 930. (A) Overall genetic load evolution. The load due to variants that are fixed in all populations is not included. (B) Difference in load between the two populations. (C) Differences in additive load. The dashed lines represent the asymptotic results from Equations (8) and (10).

**Figure S5.**
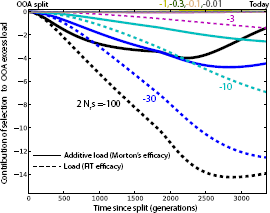
Integrated contributions of selection to load differentiation according to FIT (total load) and Morton (additive load) for partially recessive alleles (*h* = 0.2). Both show an overall increased effect of selection in the OOA population. However, genetic load remains higher in the OOA population because of the contribution of drift (Figure S3).

**Figure S6.**
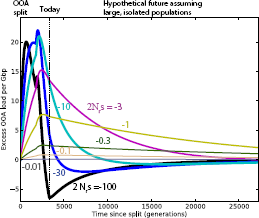
Excess recessive load in OOA populations, assuming continued future isolation between large populations (*N* = 20*N_r_*). This serves to illustrate that the ‘purging’ effect of the bottleneck on deleterious variants is observed for all alleles with *N_r_s* < –3, but that it would require much more time to compensate for the initial loss in fitness for mildly deleterious alleles with *N_r_s* > –30.

**Figure S7.**
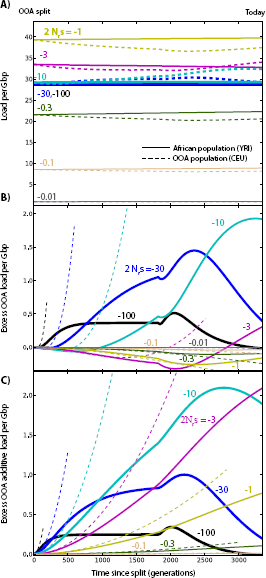
Changes in load after the OOA split model illustrated in Figure 3A, assuming dominant deleterious variants (*h* = 1), as a function of s and given ancestral population size *N_r_* = 11, 930. (A) Overall genetic load evolution. The load due to variants that are fixed in all populations is not included. (B) Difference in load between the two populations. (C) Differences in additive load. The dashed lines represent the asymptotic results from Equations (8) and (10).

**Figure S8.**
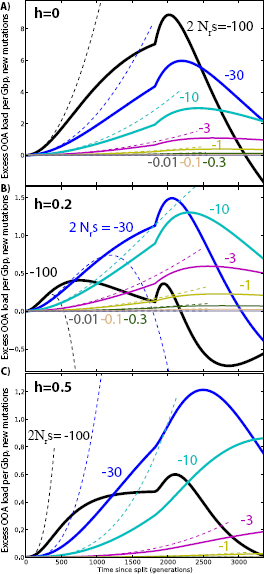
Contributions of recent mutations to load differentiation across populations, for different dominance coefficients. Here, ‘new mutations’ are mutations that occurred after the OOA split. Solid lines are the results of simulations, and dotted lines represent asymptotic values from Equation (A13).

**Figure S9.**
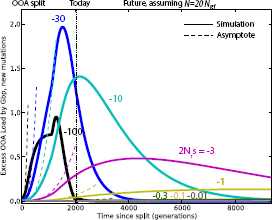
Changes in load caused by new mutations after the OOA split model illustrated in Figure 3A, given *h* = 0.5 and ancestral population size *N_r_* = 11, 930. The dashed lines represent the asymptotic results from Equation (A13).

**Figure S10.**
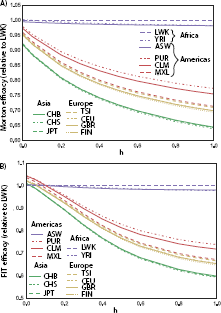
A) Present-day Morton efficacy of selection for nearly neutral variants, estimated from 1000 Genomes non-synonymous variation, assuming different dominance coefficients *h* B) FIT efficacy for the same variants.

**Figure S11.**
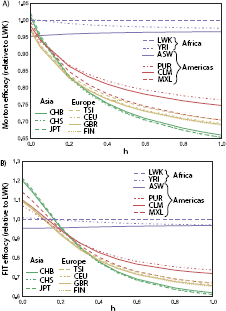
A) Present-day Morton efficacy of selection for nearly neutral variants, estimated from 1000 Genomes variants with CADD score above 2, assuming different dominance coefficients *h*. B) FIT efficacy for the same variants.

